# INPP5D limits plaque formation and glial reactivity in the APP/PS1 mouse model of Alzheimer’s disease

**DOI:** 10.1101/2022.04.29.490076

**Authors:** EL Castranio, P Hasel, J-V Haure-Mirande, AV Ramirez Jimenez, W Hamilton, RD Kim, M Wang, B Zhang, S Gandy, SA Liddelow, ME Ehrlich

**Author notes:** Equal contribution (EL Castranio, P Hasel).

## Abstract

The dual specificity lipid/protein phosphatase SHIP1 (encoded by the *INPP5D* gene) is enriched in myeloid cells. Single nucleotide polymorphisms (SNPs) in *INPP5D* coding and non-coding regions impact risk for developing late onset sporadic Alzheimer’s disease (LOAD). We present pathological analyses with spatial transcriptomics of mice with tamoxifen-sensitive microglial knockdown of *Inpp5d* and show exacerbated plaque pathology, plaque-associated microglial density, and altered gene expression around plaques, suggesting novel markers for plaque-associated reactive microglia.

## MAIN TEXT

Late-onset sporadic Alzheimer’s disease (LOAD) is the most common form of dementia, characterized by progressive memory decline. Genome-wide association studies (GWAS), whole genome sequencing, and gene-expression network analysis have identified hub and driver genes and their networks associated with risk of LOAD^1,2^. Expression of a large proportion of these genes is specific to, or enriched in, microglia, underscoring a potential role for microglia in AD pathogenesis^3^. Inositol polyphosphate-5-phosphatase D (*INPP5D*) has been highlighted by multiple approaches as an AD risk gene, and its expression in brain is restricted to microglia^4–6^. *INPP5D* encodes the 5’ phosphatase SHIP1 (Src homology 2 [SH2] domain-containing inositol polyphosphate 5’ phosphatase) and current evidence suggests that increased INPP5D levels associated with the common variant rs35349669 confer increased risk for LOAD^7,8^.

Phosphoinositides (PtdIns) are lipid molecules that exert complex effects including spatial positioning and activation of autophagic vacuoles and endosomes^9^. SHIP1 negatively regulates targets such as immune cell surface receptors by catalyzing the conversion of PtdIns(3,4,5)P3 to PtdIns(3,4) P2. This lipid dephosphorylation reaction prevents receptor activation, thereby mitigating signal transduction through the phosphatidylinositol-3-kinase/mammalian target of rapamycin (PI3K/mTOR) pathway^10^. In particular, SHIP1 inhibits phagocytic signal transduction by Triggering Receptor Expressed on Myeloid cells 2 (TREM2) and Tyrosine kinase Binding Protein (TYROBP, also known as DAP12)^11^. Constitutive deletion or pharmacologic inhibition of SHIP1 increases hematopoietic cell proliferation and phagocytosis of various substances, such as zymosan in SHIP1 knockout macrophages and amyloid beta (Aβ) in inhibited microglia^12,13^. *INPP5D* expression increases with progression of clinical LOAD^8^, but the specific role(s) that *INPP5D* play(s) in either early or late disease, and the mechanism(s), remain unknown.

Effects of alterations in the microglial phagocytic pathway on the pathogenesis of AD vary according to disease stage^14–17^. We therefore knocked down *Inpp5d* conditionally in microglia to determine early effects as the disease is developing at the preclinical level. This strategy also avoids the developmental effects that can occur in constitutive knockouts^18,19^. We used a combination of previously established mouse lines, including the PSAPP (*APP^KM670/671NL^/PSEN1^Δex-on9^*) mouse, to assess pathological consequences of *Inpp5d* knockdown (Fig. 1a). To induce recombination, 3-month-old *Inpp5d^fl/fl^/Cx3cr1^CreER/+^* mice, with and without the *PSEN1* and *APP* transgenes, were injected for 5 consecutive days with either tamoxifen (TAM) or corn oil (CO). SHIP1 protein levels were reduced by 75% one month post-recombination (Fig. 1b; Extended Data Fig. 1).

**Figure 1.**
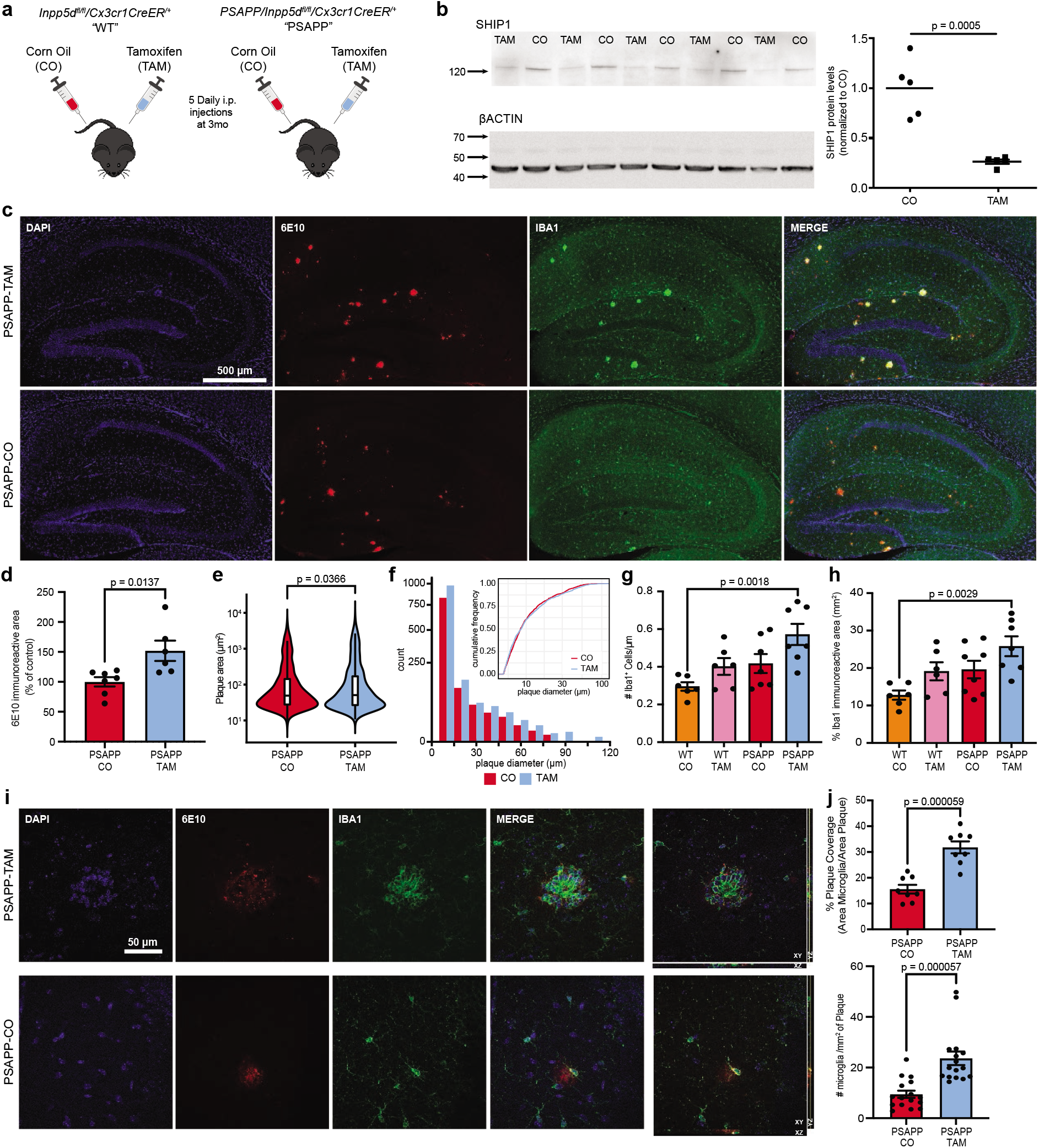
Inpp5d/SHIP1 knockdown worsens amyloid pathology and increases plaque-associated microglia number. **a)** *Inpp5d^fl/fl^/Cx3xr1^CreER+^* mice were crossed with the amyloid depositing *APP^KM670/671NL^/PSEN1^Δexon9^* mouse model of AD to obtain either heterozygous PSAPP mice, i.e. *PSAPP/Inpp5d^fl/fl^/Cx3xr1^CreER+^* (PSAPP) or their wildtype littermates *Inpp5d^fl/fl^/Cx3xr1^CreER+^* (WT). At age 3 months, male and female mice of both genotypes received either 75 mg/kg i.p. daily injections of tamoxifen (TAM) or corn oil (CO) consecutively for 5 days before aging to the experimental timepoint of ages 6-6.5months. **b)** Western blot of hippocampal tissue from WT-TAM and WT-CO at 1 month after tamoxifen-induced recombination using SHIP1 (D1163) antibody normalized to β-actin. Quantification at right. **c)** Representative images from 6-month PSAPP-TAM (top) and PSAPP-CO (bottom) mice stained for 6E10 (amyloid, red) or IBA1 (myeloid cells, green). Scale bar = 500 μm. **d)** Percentage of 6E10-immunoreactive area in the hippocampi of male and female PSAPP-CO and PSAPP-TAM mice. **e)** Quantification of 6E10-stained plaque area per plaque for PSAPP-CO and PSAPP-TAM mice. * p < 0.05. **f)** Histogram showing the size distribution (diameter, 10 μm bins) of plaques in PSAPP-CO and PSAPP-TAM mice. Insert: cumulative frequency of plaque diameter across both PSAPP-CO and PSAPP-TAM highlights relative frequency of plaque sizes across the genotypes. **g)** Quantification of number of IBA1^+^ microglia and **h)** area of IBA1^+^ area in hippocampus from WT-CO, WT-TAM, PSAPP-CO and PSAPP-TAM (n = 6-8 mice per group). **i)** Confocal Z-stack and orthogonal views of plaques from PSAPP-TAM and PSAPP-CO mice with quantification of plaque-associated microglia. N = 17-18 plaques from 5 mice per group. Scale bar = 50 μm. **j)** Quantification of plaque coverage by microglia (% area of IBA1 staining overlapping area of 6E10^+^ plaques) for PSAPP-CO and PSAPP-TAM (n = 7-8 mice per group). Bar graphs represent the mean and error bars represent SEM. P-values were calculated using unpaired t-tests for b, d, e and j, or by one-way ANOVA with Tukey’s multiple comparisons for g and h.

At 6 months of age, the percent area of 6E10 immunoreactive deposits was increased by over 50% (Fig. 1c-d) in the hippocampi of TAM-treated *PSAPP/Inpp5d^fl/fl^/Cx-3cr7^CreER/+^*(PSAPP-TAM) compared to CO-treated *PSAPPI Inpp5d^fl/fl^/Cx3cr1^CreER/+^* (PSAPP-CO) littermates. PSAPP-TAM mice also had an increase in the area and diameter of brain amyloid plaques per plaque compared to PSAPP-CO mice (Fig. 1e-f). Staining with the Congo red derivative Methoxy-XO4 (XO4), which binds specifically to β-pleated fibrils, confirmed the increase in amyloid burden associated with the *Inpp5d/SHIP1* knockdown (Extended Data Fig. 2a-b). Fibrillar deposits were increased despite unchanged levels of Aβ40, Aβ42, or Aβ42/Aβ40 ratios as determined by ELISA assays of TBS-, Triton-X-, and formic acid extracts of mouse brain homogenates (Extended Data Fig. 3).

As both microglia and astrocytes contribute to the immune response in AD^20,21^, we next measured if *Inpp5d* knockdown altered their numbers. Microglia numbers are increased in PSAPP mice with prominent peri-plaque localization^22^. In the CO- and TAM-treated *Inpp5d^fl/fl^/Cx3cr1^CreER/+^* (WT-CO, WT-TAM), the number of IBA1^+^ cells or IBA1 immunoreactive area in the hippocampus was unchanged following *Inpp5d* depletion (Fig.1g-h, Extended Data Fig. 2e). Similarly, these parameters did not change when comparing PSAPP-CO to PSAPP-TAM animals (Fig. 1c,g-h). There was an increased number of IBA1+ cells and IBA1 immunoreactive area in PSAPP-TAM mice compared to WT-CO mice (Fig. 1g-h). Most striking was a spatially-constrained increase in microglia associated with Aβ-plaques (Fig. 1c,i-j). We did not observe a significant increase of GFAP^+^ cell numbers or immunoreactive area in the hippocampus in association with *Inpp5d* knockdown (Extended Data Fig. 2a,c-d).

We next performed spatial transcriptomics to analyze brain region-resolved differential gene expression due to the presence of the APP/PS1 transgenes and the knockdown of *Inpp5d* (Fig. 2; Extended Data Figs. 4–8). We sequenced a total of 12 coronal brain sections from WT-CO, WT-TAM, PSAPP-CO and PSAPP-TAM animals (Fig. 2a) and reliably identified brain regions based on gene expression profiles (Fig. 2b,c). We found that one cluster, Cluster 26, was exclusively expressed in PSAPP mice (Fig. 2d). Cluster 26 showed a highly cluster-specific gene expression profile (Fig. 2e) and co-localized with 6E10-positive plaques of adjacent sections (Fig. 2f). Gene ontology analysis of Cluster 26 genes included “Microglia pathogen phagocytosis pathway” (e.g. include: *C1qa/b/c*, *Fcer1g*, *Vamp8*), “Tyrobp causal network” (*Tyrobp*, *Cd37*, *Cd84*, *Itgam*, *Spp1*), “Lysosome” (*Cd63*, *Hexa/b*, *Ly86*), and “Antigen processing and presentation” (*B2m*, *H2-D1/K1*, *Axl*) (Fig. 2g). Given that Aβ-plaques have been associated with increased inflammation^23^, we next asked whether previously reported reactive microglial or astrocyte sub-states have similar gene expression profiles and are present in the same spatial location as Cluster 26. Indeed, Cluster 26 has a unique gene expression profile, but we found overlapping gene sets when compared to previously reported, biologically-defined reactive glial sub-states. This included integration with DEGs from Disease Associated Microglia, both early (DAM1) and late (DAM2)^24^, Disease Associated Astrocytes (DAA)^25^, Plaque-Induced Genes (PIGs)^26^, and our previously identified reactive astrocyte interferon super-responder substate (C8_Hasel)^27^ (Fig. 2h). While DAMs and C8_Hasel reactive sub-states co-localized with Cluster 26, DAAs have a broad expression throughout the brain, which is not surprising given the many non-specific homeostatic astrocytic genes present within the module.

**Figure 2.**
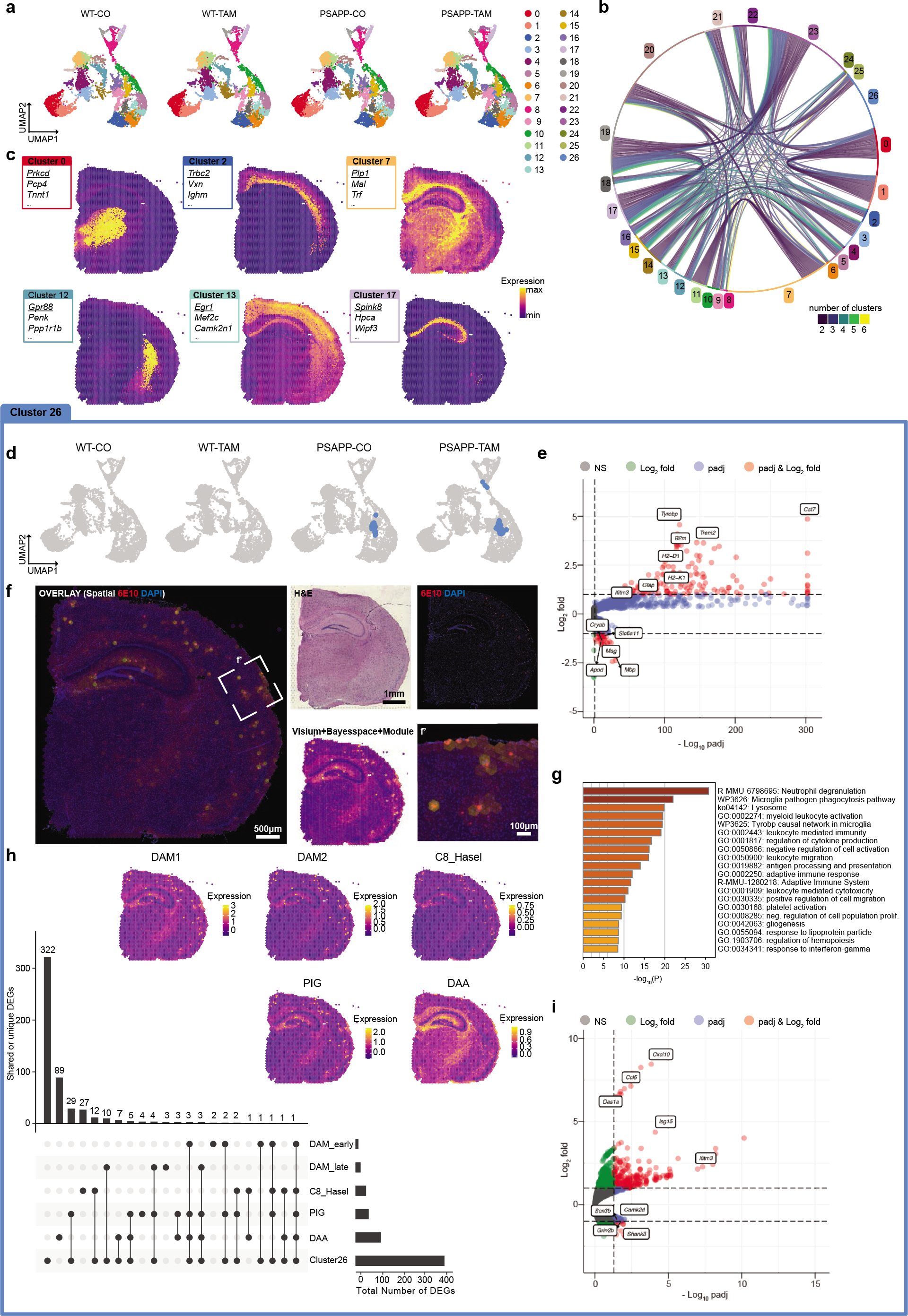
Spatial Transcriptomics identifies plaque-specific gene expression profiles exacerbated in Inpp5d/SHIP1 knockdown animals. **a)** UMAP dimensional reduction of WT-CO (8,037 spots, 3 sections), WT-TAM (8,882 spots, 3 sections), PSAPP-CO (7,748 spots, 3 sections) and PSAPP-TAM (9,366 spots, 3 sections). Each dot represents a Visium transcriptomics spot. The 26 color-coded clusters correspond to brain areas/regions (see below). **b)** Chord diagram of genes enriched >1 log_2_ fold (l2f) in each cluster and adjusted p-value <0.05. Each string represents a gene shared between 2-6 other clusters. **c)** Example clusters highlighted in a Visium brain section. Boxes show top 3 enriched genes per cluster. Underlined gene is highlighted in the SpatialFeaturePlot. **d)** UMAP with Cluster 26 highlighted showing that it is only present in PSAPP animals. **e)** Volcano plot of genes enriched in Cluster 26 as calculated with Seurat’s FindAllMarkers function. Red genes are l2f>1 and padj<0.05, blue l2f<1 and padj <0.05, green genes are l2f>1 and padj>0.05, grey genes are l2f<1 and padj >0.05. **f)** Module score of Cluster 26 and 6E10 (plaque) staining of the adjacent brain section shows colocalization. f’ shows higher magnification of box in f. **g)** GO term analysis of genes enriched l2f>1 in cluster 26 using metascape^39^. **h)** UpSet plot highlighting the overlapping and unique genes between previously defined reactive microglia and astrocyte subtypes: Disease-associated microglia, DAM1 (early) and DAM2 (late)^24^, C8_Hasel^27^, plaque-induced genes (PIGs)^26^ as well as disease associated astrocytes (DAA)^25^. Insets are module scores of the genes in each reactive sub-type highlighted on a PSAPP-TAM spatial transcriptomics brain section. For C8_Hasel, DAAs and Cluster 26, genes were filtered for padj <0.05 and l2f >0.5. For DAM1, DAM2 and PIGs, genes were selected based on gene sets provided in the original publication. **i)** Genes differentially expressed in Cluster 26 between APP CO and APP TAM as calculated with edgeR on sum of counts using muscat (red genes are l2f>1 and padj<0.05, blue l2f<1 and padj <0.05, green genes are l2f>1 and padj>0.05, grey genes are l2f<1 and padj >0.05).

We next asked whether *Inpp5d* knockdown alters periplaque gene expression patterns. We employed muscat^28^, a cluster-resolved, pseudobulk-based DEG analysis approach for multi-sample, multi-condition experiments. We used edgeR within muscat to calculate genes differentially expressed between PSAPP-CO and PSAPP-TAM brains. We found that the gene expression profile in Cluster 26 was altered by the knockdown of *Inpp5d* (Fig 2i). Of the 231 genes induced in Cluster 26 upon knock down of *Inpp5d* (padj <0.05, l2f >1), 126 were specific to PSAPP-TAM plaques. GO term analysis of these genes include an enrichment for “Defense response to virus” (*Cxcl10*, *Ifit3*, *Oas1a*, *Ccl5*, *Oasl1*), “Type I and II Interferon signaling” (*Stat1*, *Ifit2*, *Irf5/6*, *Isg15*) and “Chemokine signaling pathway” (*Plcb2*, *Rac2*, *Ccl9/12*, *Txnip*, *H2-T23*). While we discovered most DEGs in Cluster 26 due to its plaque association (Fig. 2f, Extended Data Figs. 5–7), muscat also uncovered DEGs outside Cluster 26 when comparing WT-TAM to PSAPP-TAM as well as PSAPP-CO to PSAPP-TAM. Given the recent evidence for Cre-mediated activation of microglia^29^, we calculated genes that were changed in WT-CO versus WT-TAM mice and found that while tamoxifen injection alone drove modest numbers of DEGs, they did not fall into categories associated with IFN signaling. Lastly, we employed NICHES, a recently developed package based on the fantom5 database^30^, to interrogate which putative receptor-ligand interactions were enriched within Cluster 26 and therefore plaques. We confirm that NICHES reliably detects receptor-ligand pairs along anatomical regions. In Cluster 26, we observed receptor-ligand pairs primarily involved in phagocytosis and lysosome activity, including (ligand – receptor): C1qb – Lrp1, C3 – Lrp1, Timp1 – Cd63, Pros1 – Axl and Ly86 – Cd180 (Extended Data Fig. 8).

To expand our NICHES receptor-ligand interaction spatial transcriptomic maps, we employed network analysis on clus-ter-specific DEG lists. Leveraging high-throughput molecular profiling techniques, we generated a cohort of matched whole-genome sequencing (WGS) and RNA-seq data across 4 brain regions, from a set of 364 well-characterized AD and control brains in the Mount Sinai Brain Bank (MSBB) cohort^31^. Using Bayesian probabilistic causal network (BN) analysis^32–34^, we built integrative network models to organize genome-wide gene expression features into regulatory networks^31^. When projecting the Cluster 26 expression signature and cluster-specific DEG list onto the network neighborhood of *INPP5D* in the parahippocampal gyrus region BN, we found that both were significantly enriched in the subnetwork within a path length of 6 of *INPP5D* (up to 16-fold enrichment, adjusted p-value = 3.1E-30; Fig. 3a). DAM genes like *Gpnmb*, *Trem2*, and *Tyrobp* were among the DEGs present in the network neighborhood of *INPP5D*. Fig. 3a displays the expression changes in human AD brain and in *Inpp5d* knockdown mice within the subnetwork of up to 4-steps from *INPP5D*. In this subnetwork, about half of the genes (59/119) were detected in the mouse study, among which 18 were significantly regulated by *Inpp5d* (19.9-fold, adjusted p-value = 3.3E-18). We therefore argue that Cluster 26 as well as *Inpp5d* knockdown signatures in PSAPP mice mirrored that observed in human AD gene networks. Indeed, when probing for human AD nodes from the *INPP5D* network, we find that these are regulated by the knock-down of *Inpp5d* and co-localize with the plaque-associated Cluster 26 (Fig. 3b).

**Figure 3.**
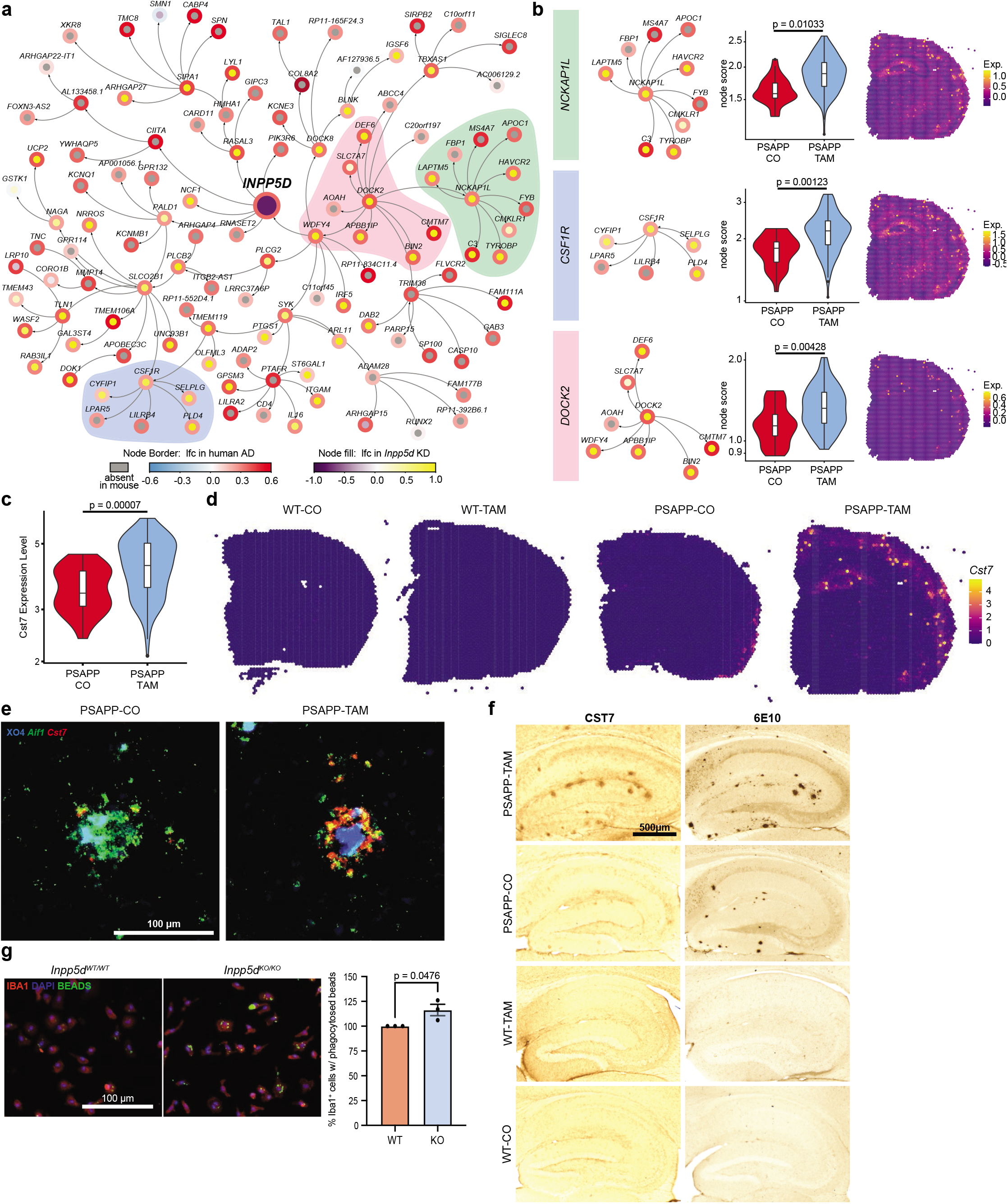
Top Cluster 26 DEG Cst7 is a plaque-specific marker and is exacerbated by *Inpp5d* knockdown. **a)** Network showing the gene expression change in human AD and *Inpp5d* knockdown mice within the neighborhood of 4 steps of *INPP5D*. Node border paint denotes the log_2_ fold change (lfc) in human AD brains compared to control brains. Node fill paint denotes the lfc caused by *Inpp5d* in mice, with grey color denoting absence in mouse data. **b)** Highlight of specific nodes from network in a (left). Genes associated with each node were enriched in cluster 26 from PSAPP-TAM compared to PSAPP-CO mice (violin plots, middle, p-values from unpaired t test) and can be spatially resolved by probing for this module in spatial transcriptomic brain sections (right). **c)** Violin plots of *Cst7* in Cluster 26 across PSAPP-CO and PSAPP-TAM shows increased expression in PSAPP-TAM plaque spots as calculated with edgeR on sum of counts using muscat. **d)** Cluster 26 marker gene *Cst7* highlighted in WT-CO, WT-TAM, PSAPP-CO and PSAPP-TAM brain sections. *e)* RNAScope of *Cst7* (red) and *Aif1* (green) with Methoxy-XO4 (blue) staining in PSAPP-CO and PSAPP-TAM mouse cortex. Scale bar = 100 μm. **f)** CST7 antibody staining detected with DAB and 6E10 in adjacent section from WT-CO, WT-TAM, PSAPP-CO and PSAPP-TAM hippocampus. Scale bar = 500 μm. **g)** Representative images of latex bead phagocytosis assay on Inpp5d-WT mouse primary microglia and Inpp5d-KO primary microglia with quantification of percentage IBA1+ cells with phagocytosed fluorescent bead (right). Scale bar = 100 μm. P-values calculated with unpaired t test.

We next probed for the gene *Cst7* (cystatin F), which is specific to Cluster 26 and which has previously been identified in DAMs and PIGs (Figs. 2e,h, 3c-d)^24,26^. In humans, *CST7* is expressed not only in microglia and macrophages but also in endothelial cells^35^. *Cst7* remained upregulated in both PSAPP-CO and PSAPP-TAM mice, suggesting that CST7 might be an excellent marker of plaques. To localize *Cst7* to specific CNS cell types, we used RNAScope in situ hybridization (Fig. 3e). *Cst7* co-localized with *Aif1*^+^ microglia and with Aβ plaques in PSAPP, but not in plaque-less WT animals. This co-localization led us to propose that Cst7 may be a novel marker for Aβ plaques and confirm that the Cst7 protein is exclusively present in PSAPP brains using IHC (Fig. 3f). Previous studies associated altered *Cst7* expression with microglial dysfunction leading to impairment of Aβ clearance^36^, and upregulation of *Cst7* in Aβ-containing microglia (but not Aβ-deficient microglia) from the 5xFAD mouse model of AD^37^. These data imply that loss of SHIP1 function after initiation of AD amyloid pathology leads to an increase in plaque burden and plaque-associated microglial gene expression.

To further assess this in light of our data suggesting an increase in plaque size (Fig. 1e-f), we performed a phagocytosis assay in vitro to compare microglia from *Inpp5d*-WT to *Inpp5d*-KO mice. We found that phagocytosis was increased in microglia that lacked *Inpp5d* (Fig. 3g), in line with previous reports in the literature^12^. We hypothesize that under chronic stimulation from Aβ in the mouse model, microglia that lack *Inpp5d* undergo functional impairment in association with Aβ-plaque deposition as was described previously^38^, resulting in more and larger plaques.

These results demonstrate that conditional *Inpp5d* downregulation in the PSAPP mouse increases plaque burden and phagocytic microglial recruitment to plaques. Our spatial transcriptomics analysis highlights an extended DEG signature associated with plaques, and identifies Cst7 as a potentially highly specific marker of plaques in the AD brain. We also take advantage of cross-species gene regulatory network analysis to highlight crossover of the *INPP5D*/SHIP1 pathway, as some aspects of plaque pathology can be species-specific. Our network analysis and extensive spatial transcriptomics data should prove helpful for other groups seeking to localize key genes of interest in the PSAPP mouse brain for their own research. It remains to be seen whether the increased plaque burden and alterations in microglia phagocytosis we describe here in PSAPP mice following *Inpp5d* knockdown also occur in other mouse models of AD (e.g. 5xFAD), or at different stages during the progression of disease pathology. Further studies will be required to investigate the mechanisms underlying SHIP1-mediated phagocytosis and plaque deposition changes in the context of AD.

## METHODS

### Animals

All animal procedures were conducted in accordance with the National Institutes of Health Guidelines for Animal Research and were approved by the Institutional Animal Care and Use Committee (IACUC) at the Icahn School of Medicine at Mount Sinai. Mice were housed on a 12 hour light/ dark cycle and had access to food and water ad libitum. *AP-P^KM670/671NL^* x *PSEN1^Δexon9^* (PSAPP), *B6.129S6-Inpp5d^tm1Wgk/J^* (*Inpp5d^fl/fl^*; stock no.: 028255), B6.129P2(Cg)-*Cx3cr1^tm2.1^*(cre/ ERT2)Litt/WganJ (*Cx3cr1^CreER/CreER^*; stock no.: 021160) and B6.C-Tg(CMV-cre)1Cgn/J (CMV-Cre; stock no.: 006054) were purchased from The Jackson Laboratory. The inducible *Cx3cr1^CreER/CreER^* were crossed with mice carrying loxP-flanked *Inpp5d* to obtain *Inpp5d^fl/fl^/Cx3cr1^CreER/+^* mice^40,41^, in which tamoxifen induced *Cre* expression results in chronic recombination in microglia, but not in transient myeloid cells. These *Inpp5d^fl/fl^/Cx3cr1^CreER/+^* mice were crossed with PSAPP mice to obtain *PSAPP/Inpp5d^fl/fl^/Cx3cr1^CreER/+^* and their wildtype, *Inpp5d^fl/fl^/Cx3cr1^CreER/+^* littermates^42^. Constitutive *Inpp5d* knockout mice were generated by crossing *Inpp5d^fl/fl^* animals to a germline Cre strain, CMV-Cre, then backcrossed to remove the CMV-Cre to obtain only *Inpp5d^KO/WT^* mice. All mice were on the C57BL/6J background and experimental groups consisted of both sexes, unless otherwise noted. Spatial transcriptomic analysis was completed on brains from females.

### Tamoxifen preparation and treatment

Tamoxifen was dissolved in corn oil at a concentration of 20 mg/ml by shaking overnight at 37 °C. For induction of Cre recombinase, 3-month-old male and female mice received daily intraperitoneal (i.p.) injections of either tamoxifen (TAM) at 75 mg/kg body weight or corn oil (CO) for 5 days.

### Tissue processing

Mice were anesthetized at 6 months of age with pentobarbital (30 mg/kg, i.p.) and transcardially perfused with 20 mL of ice-cold 0.1M phosphate-buffered saline (PBS), pH 7.4. One hemisphere was immersion-fixed for 48 hours in 4% paraformaldehyde for immunohistochemical analysis. The remaining hemisphere was dissected for isolation of hippocampus and cortex and snap-frozen for biochemical analysis. Immunohistochemistry.

Fixed hemibrains were sectioned sagittally using a vibratome (Leica) at 30 μm. Free-floating brain sections stained with 6E10 anti-Aβ antibody were pre-treated with 70% formic acid for 15 minutes. Sections were blocked for 1 hour in PBS containing 0.2% (v/v) Triton-X and 5% normal goat (or donkey) serum (NGS), then incubated overnight at 4 °C in primary antibody in PBS-0.2% Triton-X with 1% NGS. The tissues were subsequently washed in PBS-0.2% Triton-X, then incubated in PBS with 1% NGS and Alexafluor-conjugated secondary antibodies for 2 hours. Sections were then counterstained as necessary and mounted onto slides. For methoxy-XO4 staining, mounted sections were dried at RT, then incubated sequentially (5 minutes per incubation) in PBS, 40% EtOH, 1 μM methoxy-XO4 (Cayman Chemical) in 40% EtOH, and then in 90% EtOH before coverslipping. To perform DAB staining for CST7, sections were washed briefly in PBS, then underwent antigen retrieval using 1x Reveal Decloaker (Biocare Medical) at 80 °C for 15 minutes before being allowed to cool for 5 minutes. Sections were washed again in PBS before quenching with 3% H_2_O_2_, followed by permeabilization with PBS-0.2% Triton-X. Sections were blocked with 5% NGS in PBS-0.2% Triton-X for 1 hour, then incubated overnight at 4 °C in primary antibody in PBS-0.2% Triton-X with 1% NGS. The tissues were subsequently washed in PBS-0.2% Triton-X, then incubated in PBS with 1% NGS and biotinylated secondary antibody. Sections were washed in PBS, then the signal was amplified using the ABC system (VECTASTAIN Elite ABC kit, Vector Laboratories), as per the manufacturer’s instructions. The signal was then developed with a DAB Reagent (KPL), as per the manufacturer’s instructions. Sections were then mounted, dried, and then dehydrated before coverslipping and imaging. Adjacent sections were used to perform 6E10 immunostaining with DAB detection, with the sections quenched as before, then treated with 70% formic acid for 15 minutes. All steps were then performed as listed previously starting from permeabilization. The tissues were stained with the following primary antibodies: rabbit anti-IBA1 (1:500; Cat. # 019-19741, Wako), mouse 6E10 anti-Aβ (1:1000; Cat. #9320-500, Covance), chicken anti-GFAP (1:1000; NBP1-05198), and mouse anti-Cst7 (1:200; antibody F010; a gift from Dr. Adriano Aguzzi). Secondary antibodies included: goat anti-mouse IgG AlexaFluor 594 or 350 (1:1000; Invitrogen), goat anti-rabbit IgG AlexaFluor 488 (1:1000; Invitrogen), goat anti-chicken 594 (1:1000; Invitrogen); and biotinylated goat anti-mouse (1:1000, Vector Laboratories). Images were acquired on a Biotek Cytation 5 Multimode Reader (Agilent) with the same gains and exposure across all animals for each stain. Automated quantification of areas and cell counts was performed using the Biotek Gen5 software (Agilent) by creating an object mask with thresholds set for intensity and object size. For total microglia counts, adjustments were performed for plaque-associated microglia based on DAPI number. Confocal images were acquired using a Confocal Zeiss LSM 510. Images were converted to Z Projects in Fiji (NIH) based on sum intensity. Plaque-associated IBA1^+^ microglia were manually counted from Z Projects based on contact with a 6E10+ immunoreactive deposit.

### Western blot

Protein was extracted from cortical tissue by homogenization in RIPA buffer (Thermo Scientific) containing phosphatase/protease inhibitor cocktail (Thermo Scientific) and centrifuged at 15,000 xg for 20 minutes at 4 °C before collecting the supernatant. A 50 μg aliquot of total was separated by electrophoresis in precast 4-12% Bis-Tris Gels (Bio-Rad) and transferred to nitrocellulose membranes. Membranes were blocked in 5% milk before probing with the following antibodies: rabbit anti-SHIP1 (1:1000, Cell Signaling D1163) and rabbit anti-β actin (1:2500, Sigma, A-2066). The secondary antibody used was peroxidase-labelled anti-rabbit IgG (1:5000, Vector Laboratories). SuperSignal West Femto ECL (Pierce, #34096) was used to reveal the immunoreactive proteins, and images were acquired using a Fujifilm ImageReader LAS-4000. Luminescent immunoreactive protein bands were quantified using Fiji software (ImageJ).

### Spatial transcriptomics

Spatial transcriptomics analysis was performed using the 10xGenomics’ Visium platform following the manufacturer’s instructions, as we have reported^27^. In short, fresh frozen brain sections from *PSAPP/Inpp5d^fl/fl^/Cx3cr1^CreER/+^* and *Inpp5d^fl/fl^/Cx3cr1^CreER/+^* mice either treated with corn oil or tamoxifen were cryosected coronally to 10 μm and mounted on Visium capture areas (n=3 for each condition, 12 sections total). Following methanol fixation at −20 °C for 30 minutes, sections were stained with H&E, imaged on a Keyence BZ-X710 using a 20x objective and tiled using BZ-X710 software. Sections were then enzymatically permeabilized for 12 minutes, and spatial barcodes and unique molecular identifiers were added to the captured poly-A mRNA. The 12 Illumina libraries were generated using Dual Index primers and sequenced on a NovaSeq6000 using a S4 200 cycle flow cell v.1.5. The sequencing output was aligned to the mouse reference genome mm10 using SpaceRanger.

The following libraries were used to analyze and visualize the data in R (4.1.0): Seurat (4.0.6), SingleCellExperiment (1.16.0), BayesSpace (1.2.1)^43^, EnhancedVolcano (1.10.0), ComplexHeatmap (2.10.), viridis (0.6.2), muscat (1.5.2)28, ggplot2 (3.3.5), dplyr (1.0.7), purrr (0.3.4), scater (1.22.0), UpSetR (1.4.0), ggraph (2.0.5), igraph (1.2.11), gtools (3.9.2), cowplot (1.1.1), ggpubr (0.4.0), NICHES (0.0.2)^30^, CIDER (0.99.0). SCTransform was applied to each sample separately before merging. For all downstream clustering analyses, variable features from all 12 samples were used. Thirty principal components and a resolution of 0.8 were used to cluster the spots across all samples. Seurat’s FindAllMarker function was used to calculate enriched genes in each cluster and muscat was used to calculate cluster-resolved differentially expressed genes across conditions using edgeR on the sum of counts per cluster. BayesSpace^43^ was used to identify sub-spot level gene expression. NICHES was employed to calculate ligand-receptor pair enrichment using pre-defined clusters and the fantom5 database^30^ Seurat’s AddModule-Score was used to visualize groups of genes in Visium brain sections.

### Network analysis

To verify if the present mouse model gene signature mirrors that in the human AD gene network, we utilized the Bayesian probabilistic causal Network (BN) we previously constructed from the parahippocampal gyrus region of the Mount Sinai Brain Bank (MSBB) AD cohort^31,44^. The cluster signatures and cluster-specific DEGs from the spatial transcriptomics analysis were intersected with the network neighborhood within a path length of up to 6 steps from *INPP5D* in the human AD BN. P value significance of the intersections at different path lengths was computed by hypergeometric test in R (v4.0.5). A sub-network up to 4-steps from INPP5D was visualized by Cytoscape (v3.9.0).

### RNA in situ hybridization

RNA in situ hybridization (ISH) was performed using the RNAscope Multiplex Fluorescent kit (ACD) as before with slight modification^45^. The same hemibrains collected for spatial transcriptomics were sectioned at 10 μm on a cryostat and mounted on glass slides. Slides were fixed in 4% paraformaldehyde for 15 minutes at 4 °C. After fixation, sections were dehydrated and stained with methoxy-XO4 by immersion in 50% EtOH for 2 minutes, 50% 5 μM methoxy-XO4 (Cayman Chemical) in 50% EtOH for 2 minutes, then 50% EtOH for 1 minute, before proceeding through 70% EtOH and two rounds of 100% EtOH for 5 minutes each. The mounted sections were quenched with H_2_O_2_ for 10 minutes then protease-permeabilized for 20 minutes at room temperature using reagents provided in the kit (Protease IV). Brain sections were then incubated for 2 hours at 40 °C with probes followed by signal amplification steps. Once the ISH assay was complete, slides were incubated with fluorophore-conjugated secondary antibody for 30 minutes, then counterstained with DAPI. The following probes were used: *Cst7-C2* (Cat. #498711-C2) and *Aif1*-C4 (Cat. #319141-C4). Aβ assays.

Tissue was processed via serial detergent fractionation with ultracentrifugation to produce TBS-soluble, Triton-X-soluble and formic-acid-soluble Aβ fractions as described46. In brief, tissue was homogenized in ice-cold TBS containing protease/phosphatase inhibitor cocktail, then ultracentrifuged at 100,000 xg for 1 hour at 4 °C. The supernatant containing the TBS-soluble fraction was collected, and the pellet was subsequently homogenized in Triton-X solution (TBS with 1% (v/v) Triton-X-100) containing protease/phosphatase inhibitor cocktail and ultracentrifuged as before. The supernatant containing the Triton-X-soluble fraction was collected, and the pellet was homogenized in 70% formic acid before the ultracentrifugation was repeated. The resulting formic acid fraction was neutralized with 1 M Tris buffer. To quantify Aβ levels, human/rat Aβ 1–40 and 1–42 ELISA kits (Wako, Cat. #294-64701 and #290-62601) were used according to the manufacturer’s instructions. Absolute concentrations of Aβ were normalized to initial tissue weight.

### Purification of primary mouse microglia and phagocytosis assay

Heterozygous *Inpp5d^KO/WT^* mice were crossed to obtain *Inpp5d^KO/KO^* (KO) and *Inpp5d^WT/WT^* mice for isolation of primary microglia. Primary microglia were isolated from cerebral cortices and hippocampus from postnatal day P0-P3 mice as described^47^ with slight modification. In brief, tissue was homogenized in ice-cold Hibernate A (BrainBits), then centrifuged at 300 xg for 5 minutes at 4 °C. The pellet was resuspended in high glucose DMEM supplemented with 10% heat-inactive FBS (Gibco), 2 mM glutamine, penicillin/streptomycin (100 U/ml and 0.1 mg/ml respectively) and seeded in poly-L-lysine pre-coated flasks. Cells were maintained at 37 °C and 5% CO_2_, during which media was changed twice a week. Microglia were collected after approximately 8-10 days by agitating the flasks at 180 rpm for 30 minutes to detach microglial cells from the astrocytic monolayer.

Green fluorescent latex beads (Sigma #L1030) were used to assess phagocytosis in the primary microglia. Beads were pre-opsonized in heat inactivated FBS (Gibco) for 1 hour at 37 °C before use^47^. Cells were submitted to serum deprivation for 1 hour prior to addition of the beads, and then fluorescent latex beans were added to the media for an additional 1 hour or 4 hours. The final concentrations for beads and FBS in serum-free DMEM were 0.01% (v/v) and 0.05% (v/v) respectively. Cultures were then washed 3 times with PBS and fixed in 4% paraformaldehyde before cells were stained against IBA1 and DAPI. Images were obtained using a Biotek Cytation 5 Multimode Reader (Agilent) to generate (at minimum) a 3×3 tiled montage of the well. Automated quantification of primary microglia number and dual-labeling was performed using the Biotek Gen5 software (Agilent) by creating an object mask with thresholds set for intensity and object size for IBA1^+^ cells with a secondary mask to count the subpopulation of dual-labelled for green fluorescent beads.

### Human brain information and experiments

For human AD gene network analysis, we used the multi-Omics data we previously generated from a set of 364 well-characterized AD and control brains spanning the full spectrum of late-onset AD-related cognitive and neuropathological disease severities represented in the MSBB AD cohort. Details about the cohort population information, generation of-Omics data (including whole-genome sequencing and RNA sequencing), and data preprocessing are described in Wang et al^44^.

### Statistical analyses

All data, excluding spatial transcriptomics results, were analyzed with GraphPad Prism 9 and are presented as means ± SEM. Sample sizes and statistical tests are indicated in figure legends. Analyses used include one-way ANOVA, and student’s t-test.

## Acknowledgements

The authors would like to thank Dr. Adriano Aguzzi for generously providing us with the anti-CST7 antibody F010. The authors were supported as follows: E.L.C. (P30 AG066514 to Mary Sano with Developmental Pilot Award), S.G. (U01AG046170, RF1AG058469, RF1AG059319, R01AG061894, and P30 AG066514 to Mary Sano); M.E.E. (U01AG046170); M.W. (U01AG046170, RF1AG057440); B.Z. (U01AG046170, RF1AG057440); S.A.L. (NYU School of Medicine, the Blas Frangione Foundation, the Neurodegenerative Diseases Consortium from MD Anderson, Alzheimer’s Research UK, the Gifford Family Neuroimmune Consortium as part of the Cure Alzheimer’s Fund, The Alzheimer’s Association, the Alzheimer’s Disease Resource Center at NYU Langone Medical Center. He also acknowledges the generous anonymous donors, and financial support of Paul Slavik). P. H. and S. A. L. acknowledge the Genome Technology Center at NYU Langone Medical Center for sequence alignment and data pre-processing.

## Declarations of Interest

SAL is a founder of AstronauTx Ltd. All other others declare no competing interests.

---

**Extended Data Fig. 1.**
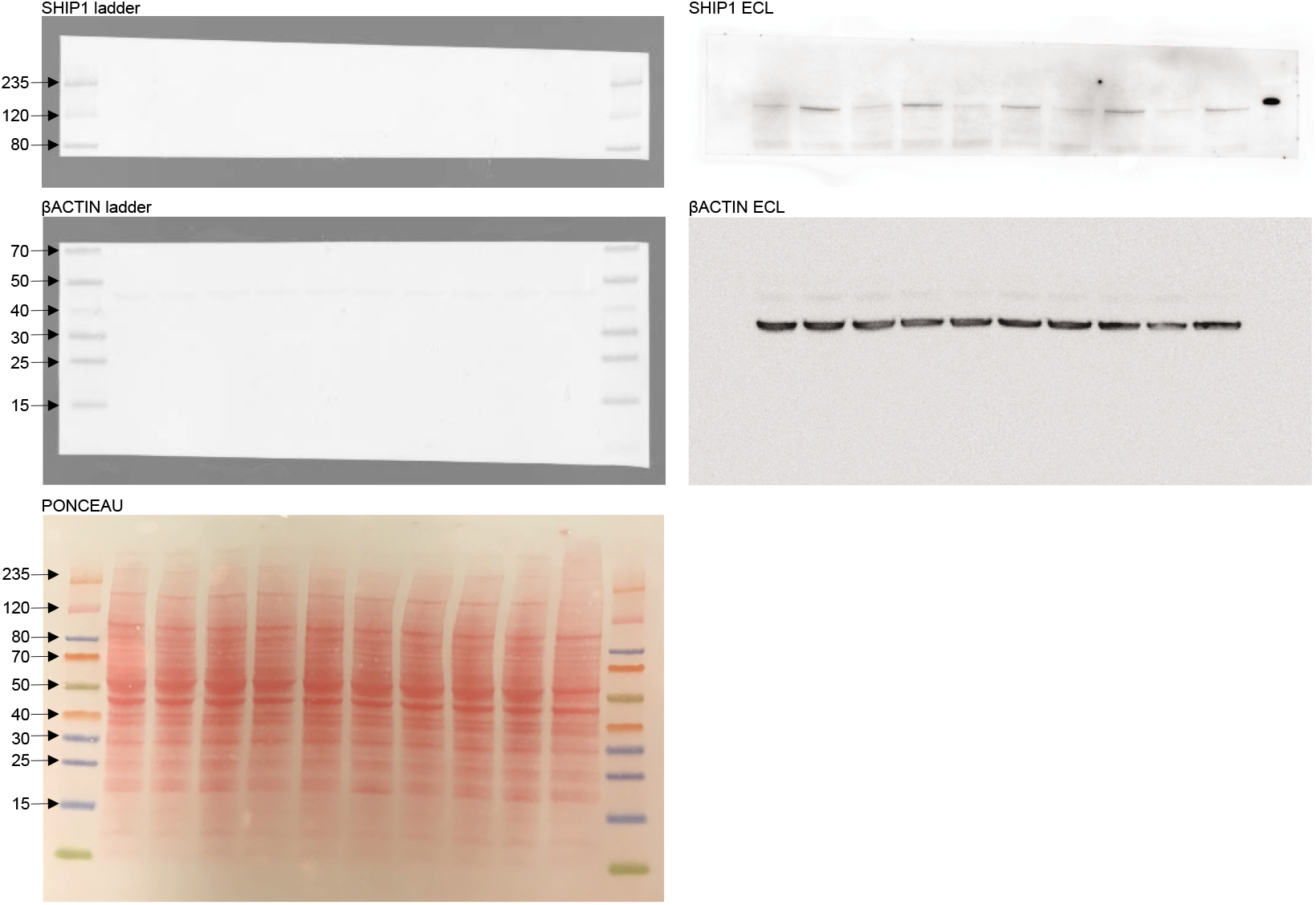
Full western blot images for SHIP1 and β-actin and Ponceau associated with Fig. 1b.

**Extended Data Fig. 2.**
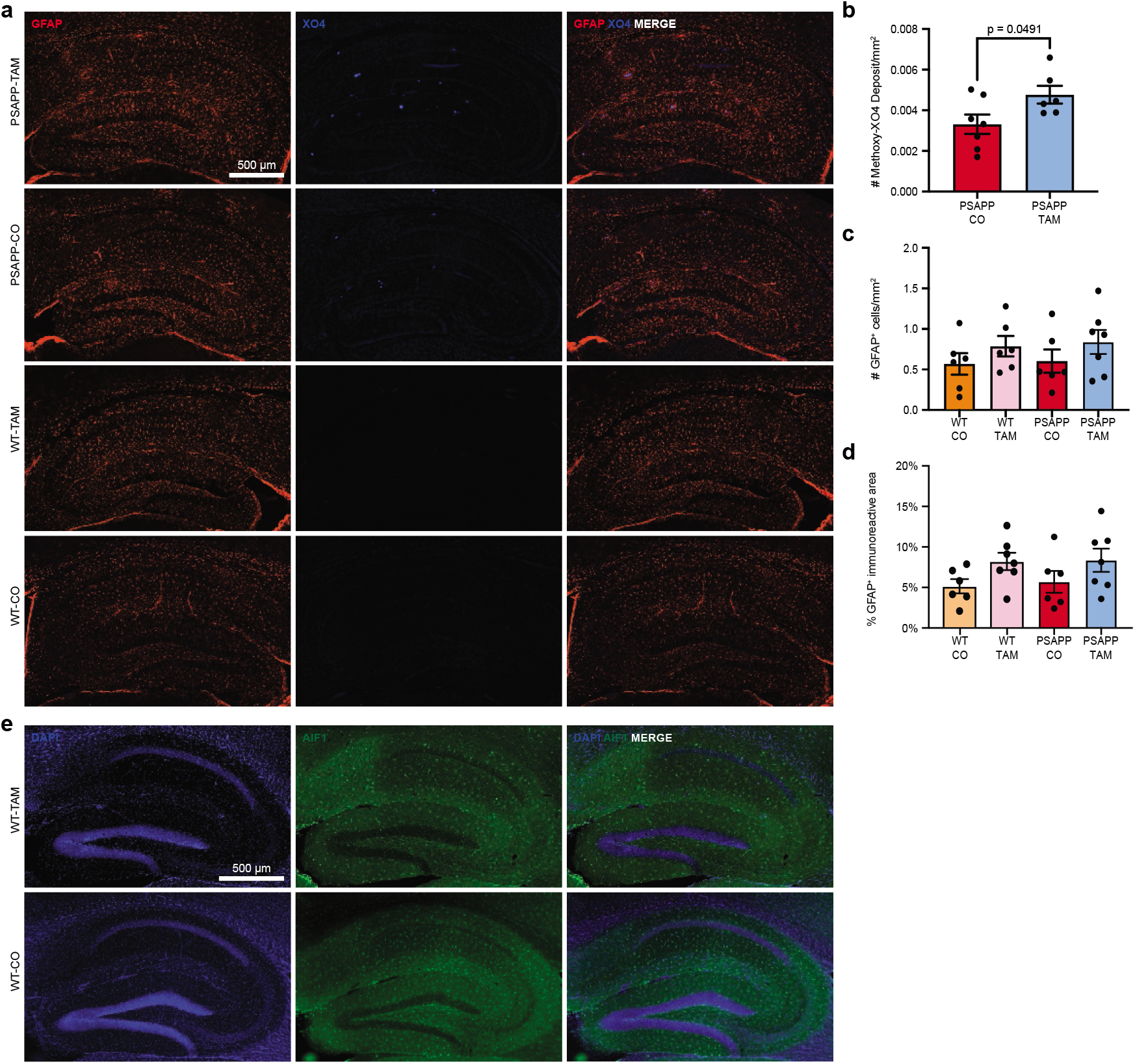
**a)** Representative images from 6 month old PSAPP-TAM, PSAPP-CO, WT-TAM and WT-CO mice stained with anti-GFAP (astrocytes, red) or XO4 (plaques, blue). Scale bar = 500 βm. **b)** Number of XO4-immunoreactive deposits in the hippocampus of male and female PSAPP-TAM and PSAPP-CO mice. **c)** Quantification of number of GFAP^+^ astrocytes and **d)** area of GFAP^+^ immunostaining in hippocampus from PSAPP-TAM, PSAPP-CO, WT-TAM and WT-CO mice (n = 7-8 mice per group). **e)** Representative images from 6 month old WT-TAM and WT-CO mice immunostained with 6E10 (amyloid, red) or anti-IBA1 (myeloid cells, green). Scale bar = 500 μm.

**Extended Data Fig. 3.**
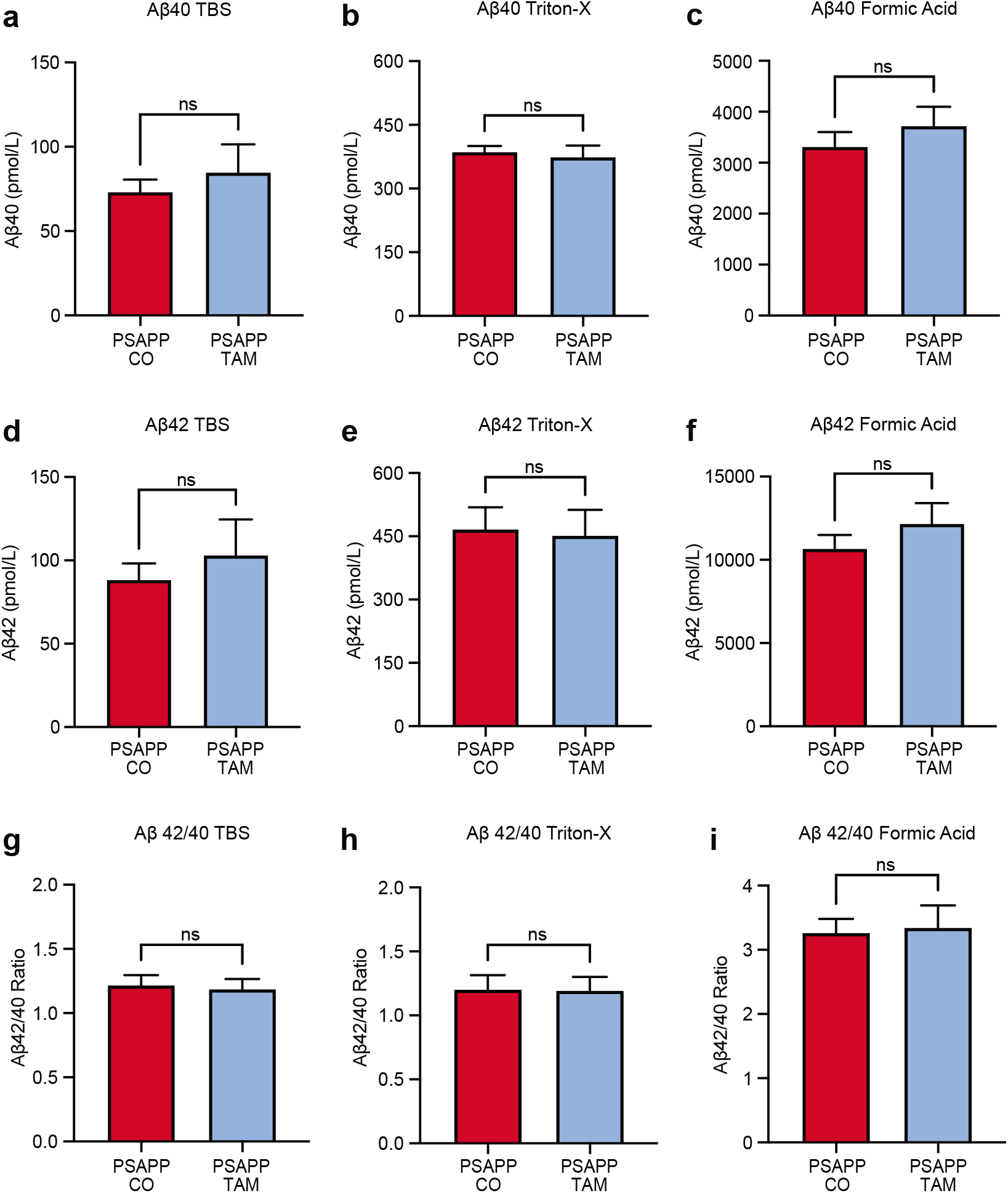
Tissues from hippocampus and posterior cortex from female PSAPP-CO and PSAAP-TAM mice were processed via serial extraction to produce fractions of TBS-, Triton-X and formic acid soluble Aβ. Levels of Aβ40 **(a-c)** and Aβ42 **(d-f)** were determined from each fraction via ELISA. The Aβ42/40 ratio was calculated for each fraction **(g-i)**. n=6 per group. ns: not significant. Data presented as mean ± SEM.

**Extended Data Fig. 4.**
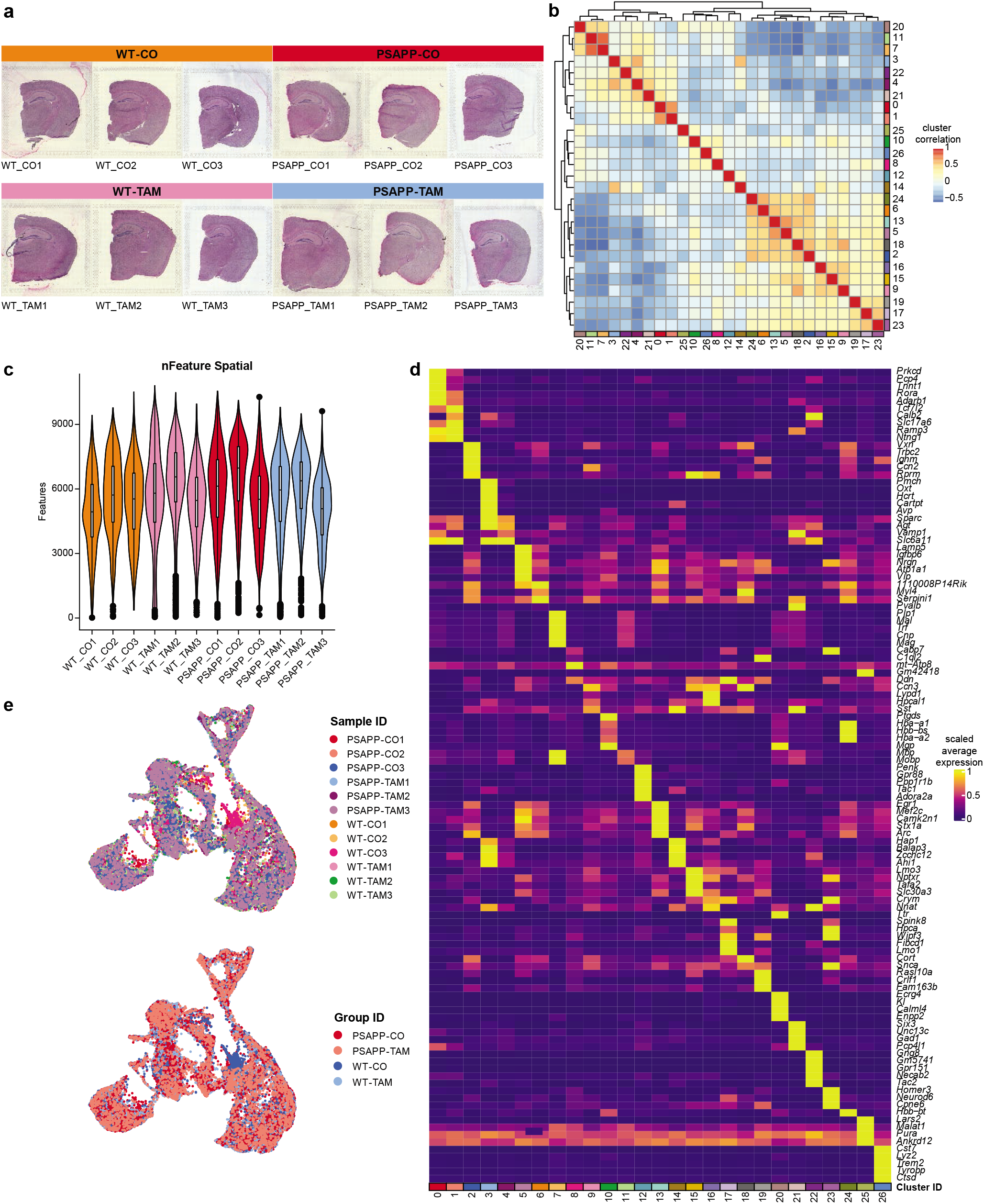
**a)** Hematoxylin & Eosin (H&E) stain of the 12 brain sections used for Visium spatial transcriptomics. **b)** Correlation matrix as calculated with CIDER shows correlation of transcriptomic cluster identities. **c)** Violin Plot showing the number of features per spot for each sample. **d)** Heatmap showing the average expression of the top 5 genes enriched within each cluster normalized to their maximal expression. **e)** UMAPs of the Visium spots grouped by sample ID (top) and group ID (bottom).

**Extended Data Fig. 5.**
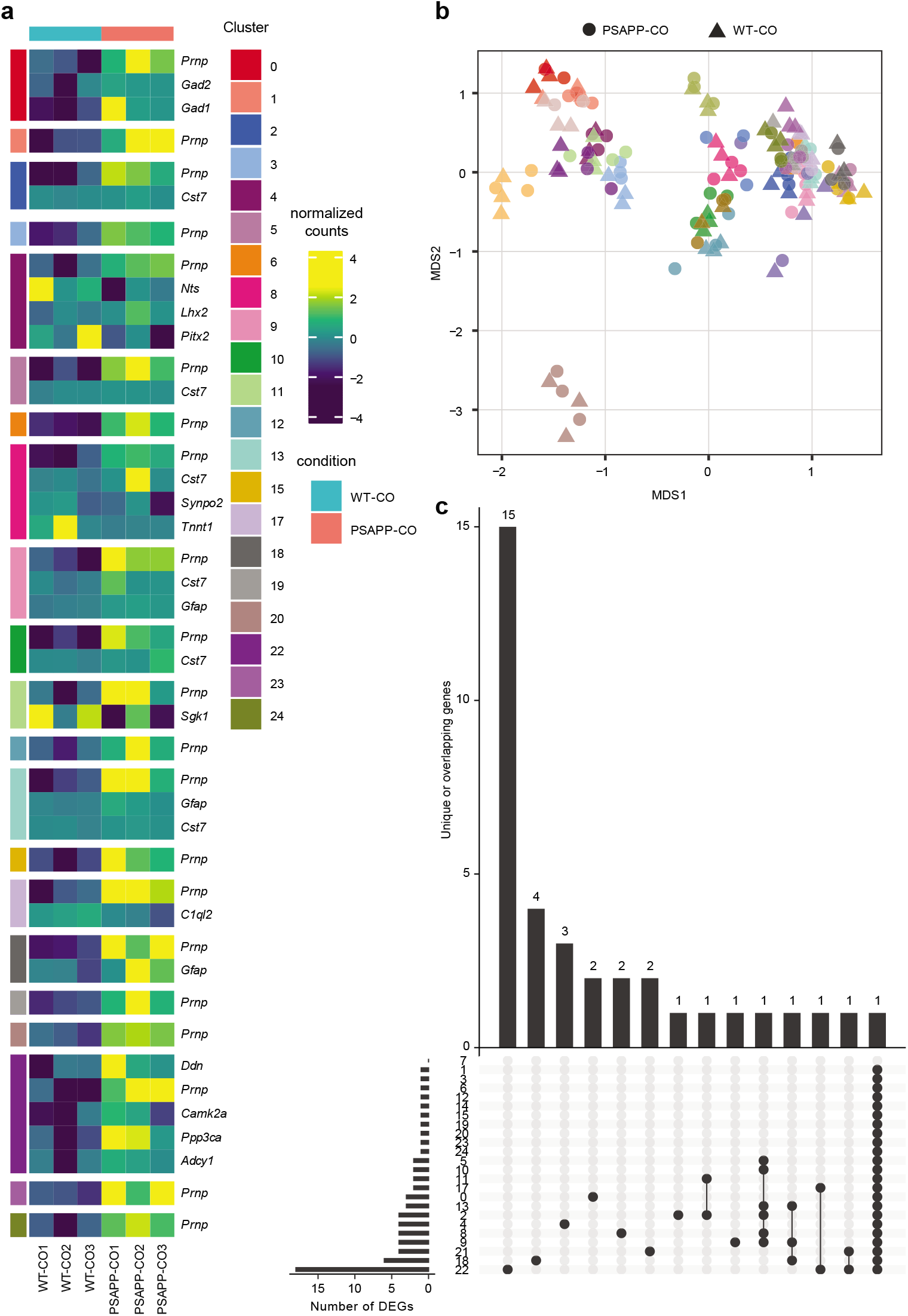
**a)** Heatmap showing cluster-resolved DEGs as calculated using edgeR on sum of counts within the muscat package when comparing WT-CO to PSAPP-CO. **b)** Multidimensional scaling (MDS) plot showing effect of PSAPP on gene expression across all clusters and all samples. **c)** UpSet plot showing unique and over-lapping DEGs with l2f > |1|. Note that Cluster 26 is only present in PSAPP mice and is therefore excluded by edgeR.

**Extended Data Fig. 6.**
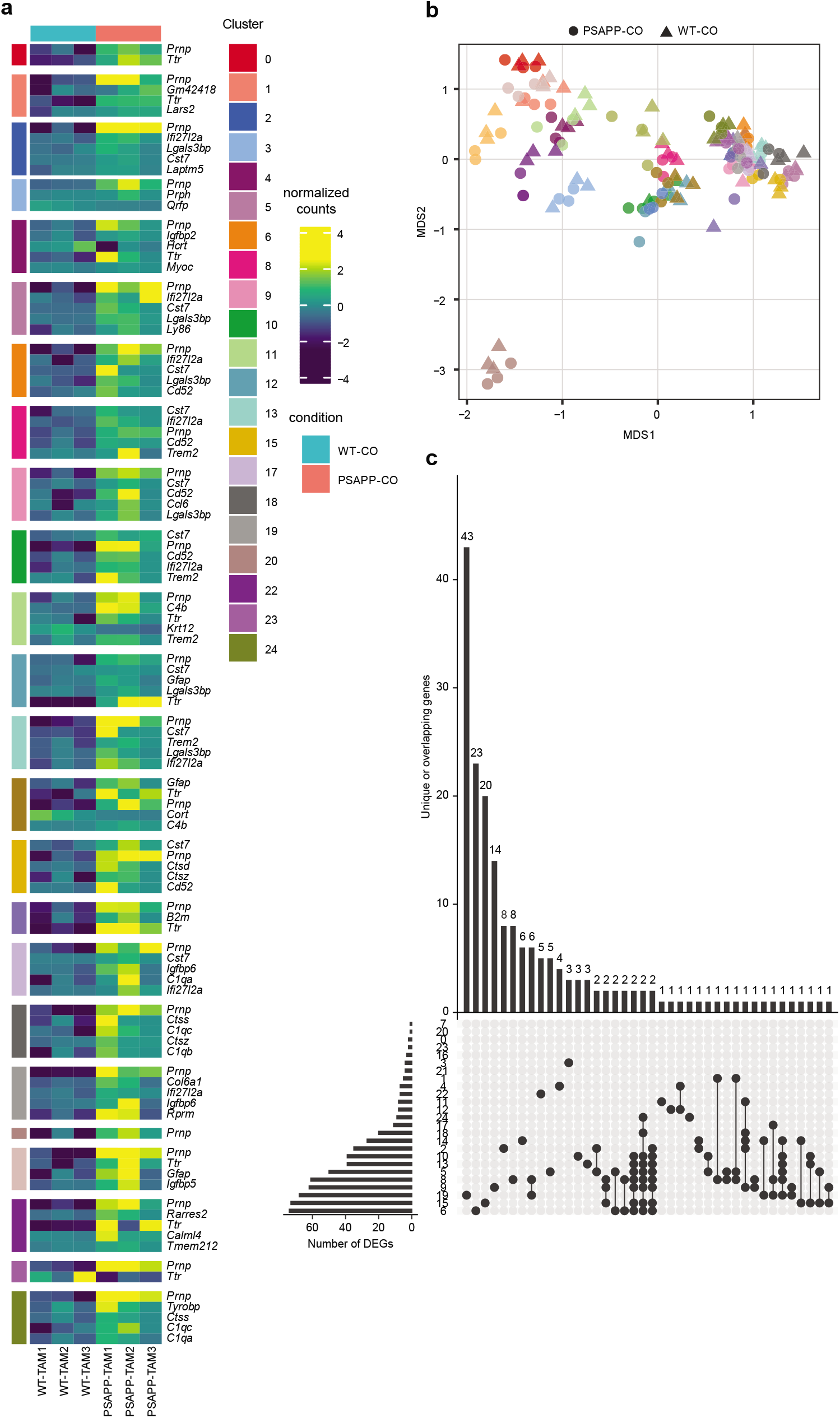
**a)** Heatmap showing cluster-resolved DEGs as calculated using edgeR on sum of counts within the muscat package when comparing WT-TAM to PSAPP-TAM. **b)** Multidimensional scaling (MDS) plot showing effect of PSAPP-TAM on gene expression across all clusters and all samples. **c)** UpSet plot showing unique and overlapping DEGs with l2f > |1|. N.B. that Cluster 26 is only present in PSAPP mice and is therefore excluded by edgeR.

**Extended Data Fig. 7.**
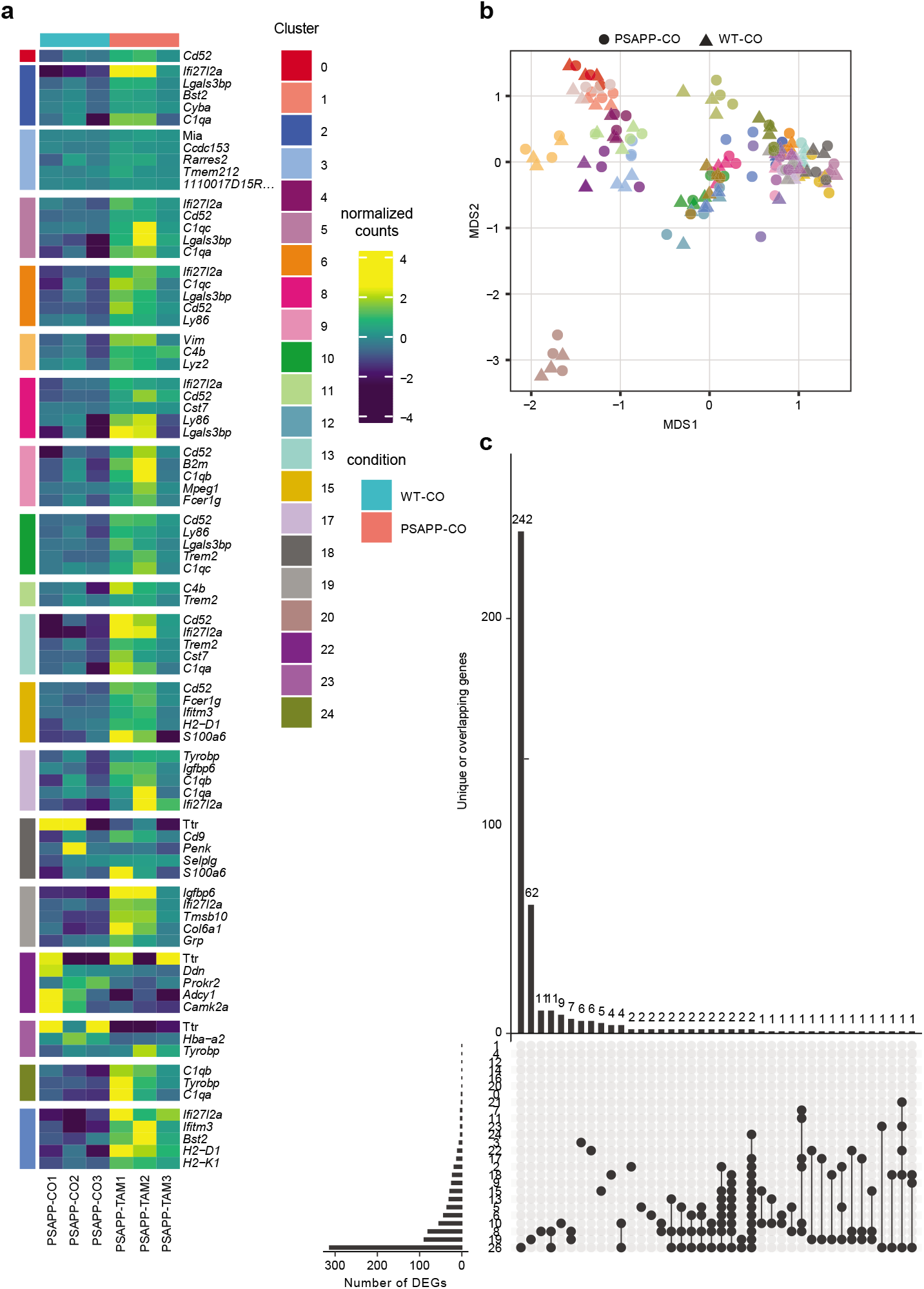
**a)** Heatmap showing cluster-resolved DEGs as calculated using edgeR on sum of counts within the muscat package when comparing PSAPP-TAM to PSAPP-CO. **b)** Multidimensional scaling (MDS) plot showing effect of PSAPP-TAM on gene expression across all clusters and all samples. **c)** UpSet plot showing unique and overlapping DEGs with l2f > |1|.

**Extended Data Fig. 8.**
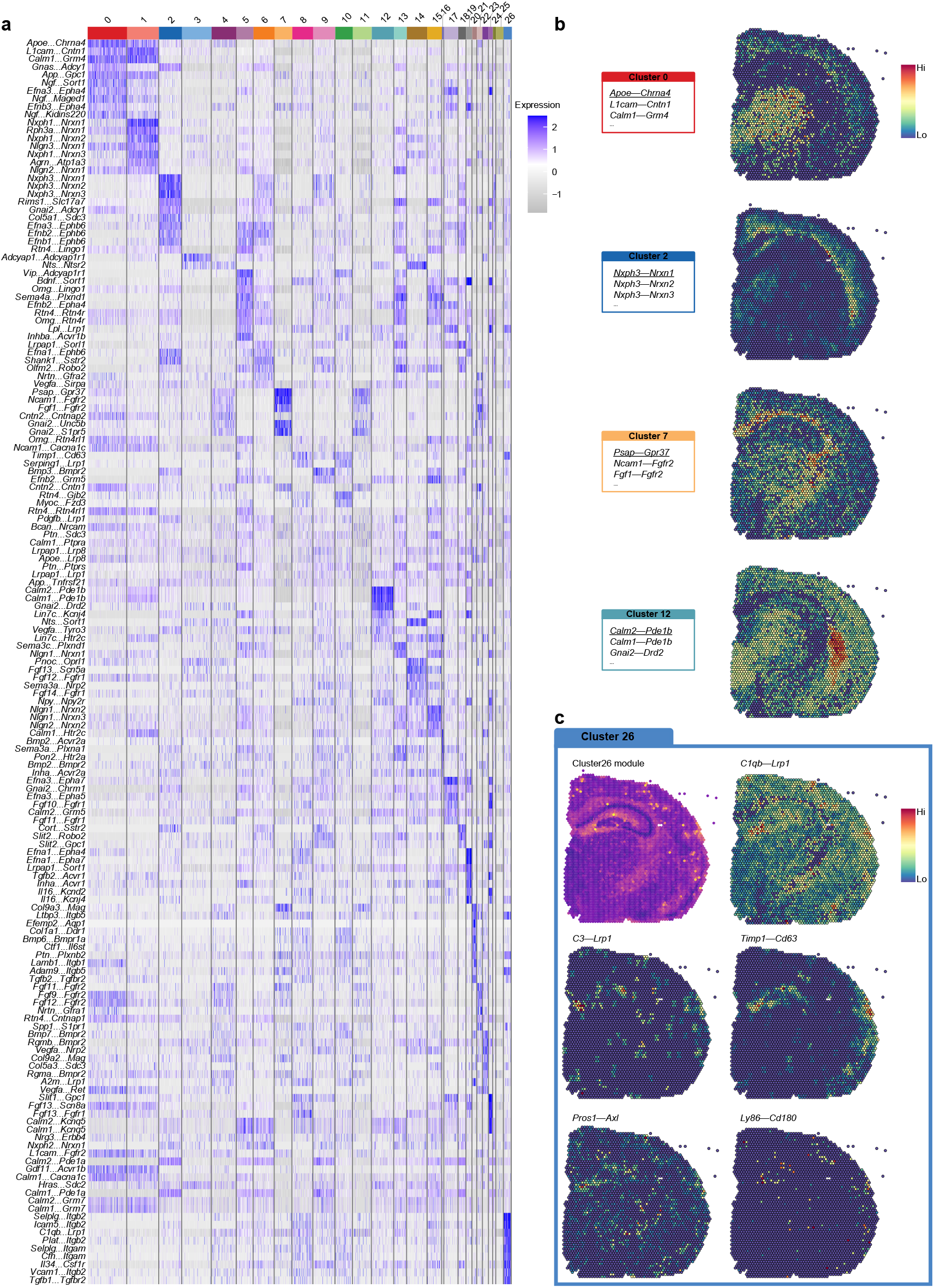
**a)** Heatmap of receptor-ligand pairs enriched in each of the 26 clusters identified using NICHES 30. b) Examples of cluster-enriched receptor-ligand pairs show receptor-ligand pairs can express along anatomical regions (see Fig. 2c for comparison). **c)** NICHES on Cluster 26 highlights enrichment of receptor-ligand pairs involved in inflammation and phagocytosis.

**Extended Data Fig. 9.**
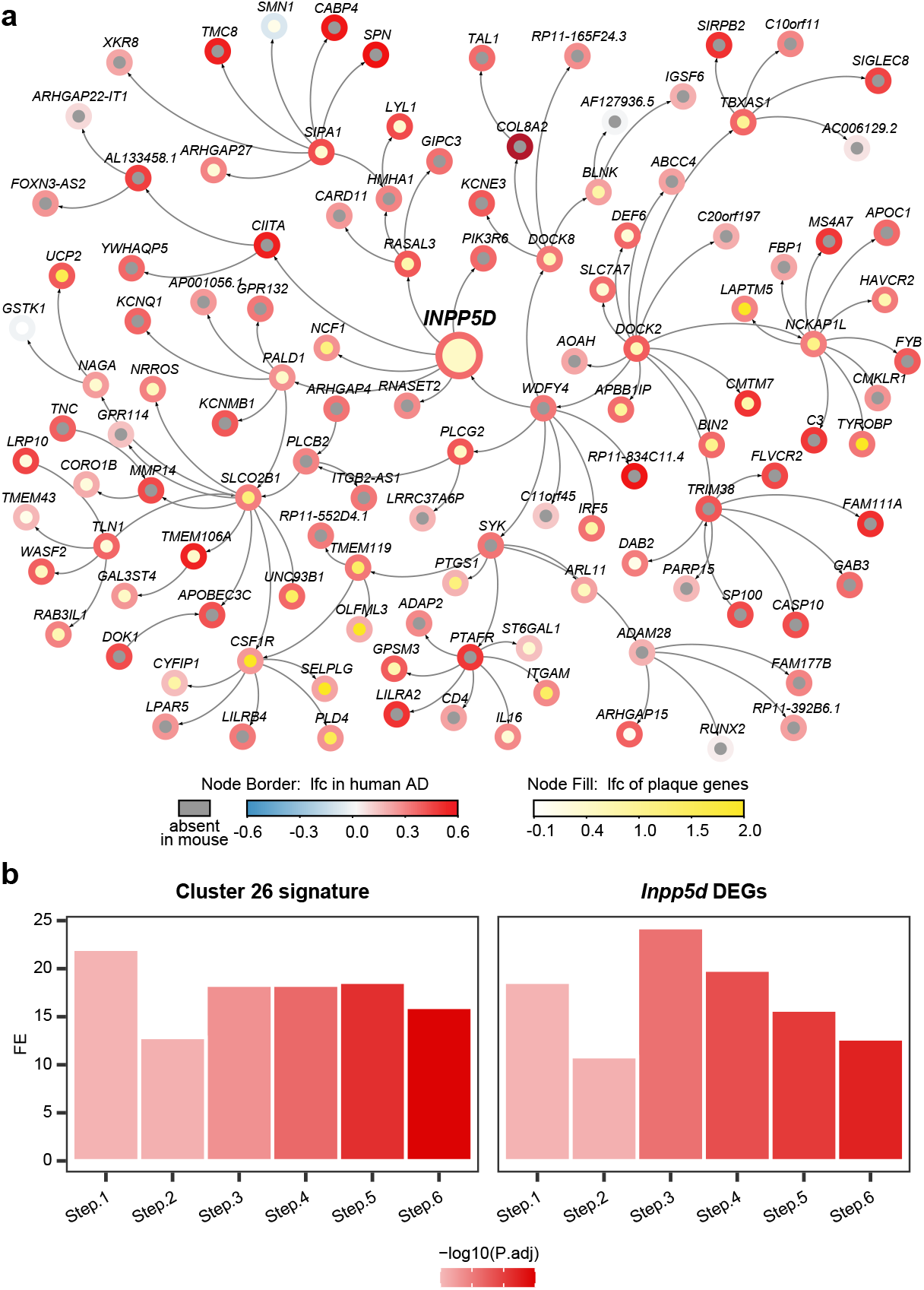
**a)** Network showing the gene expression change in human AD and Cluster 26 from *Inpp5d* knockdown mice within the neighborhood of 4 steps of *INPP5D*. Node border paint denotes the log_2_ fold change (lfc) in human AD brains compared to control brains. Node fill paint denotes the log_2_ fold change caused by *Inpp5d* in mice, with grey color denoting absence in mouse data. **b)** Barplot showing enrichment of Cluster 26 signature and Inpp5d knockdown DEGs in the network neighborhood of *INPP5D* in human AD network. X-axis denotes the neighborhood at step 1-6 from *INPP5D*. Y-axis denotes the fold enrichment (FE). Color gradient of the bars denote the P value significance of the enrichment.

